# Nano Pom-poms Prepared Highly Specific Extracellular Vesicles Expand the Detectable Cancer Biomarkers

**DOI:** 10.1101/2021.02.21.432188

**Authors:** Nan He, Sirisha Thippabhotla, Cuncong Zhong, Zachary Greenberg, Liang Xu, Ziyan Pessetto, Andrew K. Godwin, Yong Zeng, Mei He

## Abstract

Extracellular vesicles (EVs), particularly exosomes, are emerging biomarker sources. However, due to heterogeneous populations secreted from diverse cell types, mapping EV multi-omic molecular information specifically to their pathogenesis origin for cancer biomarker identification is still extraordinary challenging. Herein, we introduced a novel 3D-structured nanographene immunomagnetic particles (NanoPoms) with unique flower pom-poms morphology and photo-click chemistry for specific marker-defined capture and release of intact small EVs. This specific EV isolation approach leads to the expanded identification of targetable cancer biomarkers with enhanced specificity and sensitivity, as demonstrated by multi-omic EV analysis of bladder cancer patient tissue fluids using the next generation sequencing of somatic DNA mutations, miRNAs, and the global proteome. The NanoPoms prepared sEVs also exhibit distinctive in vivo biodistribution patterns, highlighting the highly viable and integral quality. The developed method is simple and straightforward, and is applicable to nearly all types of biological fluids and amenable for scale up and high-throughput EV isolation.

## Introduction

Despite the tremendous efforts made in developing cancer biomarkers and liquid biopsy for past decades, only a few (less than 25) cancer biomarkers have been approved by FDA for clinical practice(*1, 2*). Extracellular vesicles (EVs) have been emerging biomarker sources for expanding the landscape of cancer biomarker discovery in promoting cancer diagnosis(*3-5*), immunotherapy(*6, 7*), drug target and delivery(*8*). Significant attention has been focused on the exosome type small EVs (sEVs) and their molecular components (*e*.*g*., proteins, DNAs, mRNA and miRNA), which has been found in association with a variety of physiological functions and pathological disease states(*9*). sEV secretion is exacerbated from tumor cells and enriched with a group of tumor markers, as evidenced by increased presence in plasma and ascites from patients in variable cancers(*10*). However, currently there is no standardized purification method for obtaining pure sEV populations that are specific to their cellular origin and molecular information(*11, 12*). EVs are living cell-secreted membrane vesicles in multiple subpopulations, including membrane shedding microvesicles (100 nm-1000 nm), endosomal multivesicular body released exosomes (30 nm-150 nm), and apoptotic cellular fragment vesicles (≥ 1000 nm)(*13*). Due to such large heterogeneity and significant size overlap between vesicle populations, the consensus has not yet emerged on precisely defining EV subtypes, such as endosome derived exosomes(*14*) which is highly relevant to the disease pathogenesis. The generic term of EVs is recommended by complying with 2018 guidelines from the International Society for Extracellular Vesicles (ISEV) proposed Minimal Information for Studies of Extracellular Vesicles (“MISEV”)(*14*). Current purification methods that recover the highest amount of extracellular materials, no matter with the vesicle or non-vesicular molecules, are mainly the precipitation polymer kits and lengthy ultracentrifugation-based (UC) approach(*15, 16*). Such isolation approach is not scalable and unable to differentiate the sEV populations from different cellular origin or other EV subtypes (*e*.*g*., microvesicles and apoptotic bodies), neither free proteins(*17*) or viruses, in turn, posing a significant concern for studying cancer biomarkers from tumor cells derived sEVs. The bulk measurement of a mixture of vesicle populations could potentially mask the essential biosignatures, which severely impairs the investigations of associated pathological mechanism(*18-21*). As the perfectly enriched biomarker sources, using EVs for mapping multi-omic molecular information specifically to their pathogenesis in cancer biomarker identification is still extraordinary challenging.

Herein, we introduced a novel approach using 3D-structured nanographene immunomagnetic particles (NanoPoms), which possesses unique flower pom-poms morphology and photo-click chemistry for specific marker-defined capture and release of intact sEVs from nearly all types of biological fluids, including human blood, urine, cow’s milk, and cell culture medium, etc. Compared to current existing immunomagnetic beads based EV isolation either in small quality or bound to solid surface/particles(*22-24*), NanoPoms enable on demand capture and release of intact sEVs which leads to the expanded identification of targetable cancer biomarkers with enhanced specificity and sensitivity. The group of enriched biomarkers carried by EVs, including DNAs, RNAs and proteins, could offer the unmatched possibility to integrate multi-omic data analysis for expending the landscape of cancer biomarker discovery and precisely defining the onset and progression of cancer diseases(*25*). In this paper, we demonstrated such capability by multi-omic EV analysis of bladder cancer patient tissue fluids (e.g., urine, plasma, and tumor tissue) using the next generation sequencing (NGS) of somatic DNA mutations, miRNAs, and the global proteome, for achieving non-invasive, ultra-sensitive diagnosis of bladder cancer. The results showed much higher specificity and sensitivity for detecting urological tumor biomarkers from NanoPoms isolated sEVs compared to other ultracentrifugation or bead isolation approaches. We also identified a few new miRNAs and proteome cancer biomarkers highly enriched in urinary EVs but was not reported yet, which could expand the landscape for discovering new EV cancer biomarkers with improved diagnostic accuracy. The highly integral quality of NanoPoms prepared sEVs was demonstrated by in vivo biodistribution analysis which showed distinctive patterns from different subtypes of sEVs, implying the great potential for targeted precision therapeutic development.

## Results

### Nano pom-poms enable specific capture and on-demand release of intact sEVs

In this work, we introduce a novel 3D-structured nanographene immunomagnetic particles (NanoPoms) with unique flower pom-poms morphology and photo-click surface chemistry for specific marker-defined capture and release of intact sEVs (Fig. 1). Conventionally, the non-covalently assembled nano-graphene suffers from the instability in buffer solutions over time(*26*). Our method interfaces Fe_3_O_4_/SiO_2_ core-shell particles (∼800 nm) with graphene nanosheets via carboxamide covalent bonds, which leads to substantially improved stability in the aqueous samples. The flower pom-poms morphology produces the unique 3D nano-scale cavities in between for affinity capture of only nano-sized vesicles (Fig. 1 and Fig. S1). The dense nano-graphene hydrophilic sheet layers provide much larger surface area(*27-30*) for immobilization of affinity capture entities (e.g., antibodies, aptamers, and affinity peptides) as shown in Fig. 1b right panel in contrast to conventional beads. Most importantly, the conjugated photo-click chemistry on bead surface allows the release of intact, captured sEVs on demand, which further ensures the specificity for harvesting marker-defined sEV subpopulations. Fig. 1c TEM imaging demonstrated the capture of sEVs which exhibit much narrower size distribution than ultracentrifugation (UC) prepared EVs (Fig. 1d). We observed the dense and round-shaped sEVs (∼100 nm) completely covering the surface of NanoPoms and subsequently restoring the pom-poms surface morphology after light release (Fig. 1e). The fluorescence binding analysis was also performed to evaluate sEV capture performance and capacity (Fig. S2 and s3). This NanoPoms method is applicable to nearly all types of biological fluids, including human blood, urine, cow’s milk, and cell culture medium, etc (Fig. S3). The operation protocol is simple and cost-effective, amenable for scaling up, sterilization settings, and GMP operations (see Table s1).

**Fig 1.**
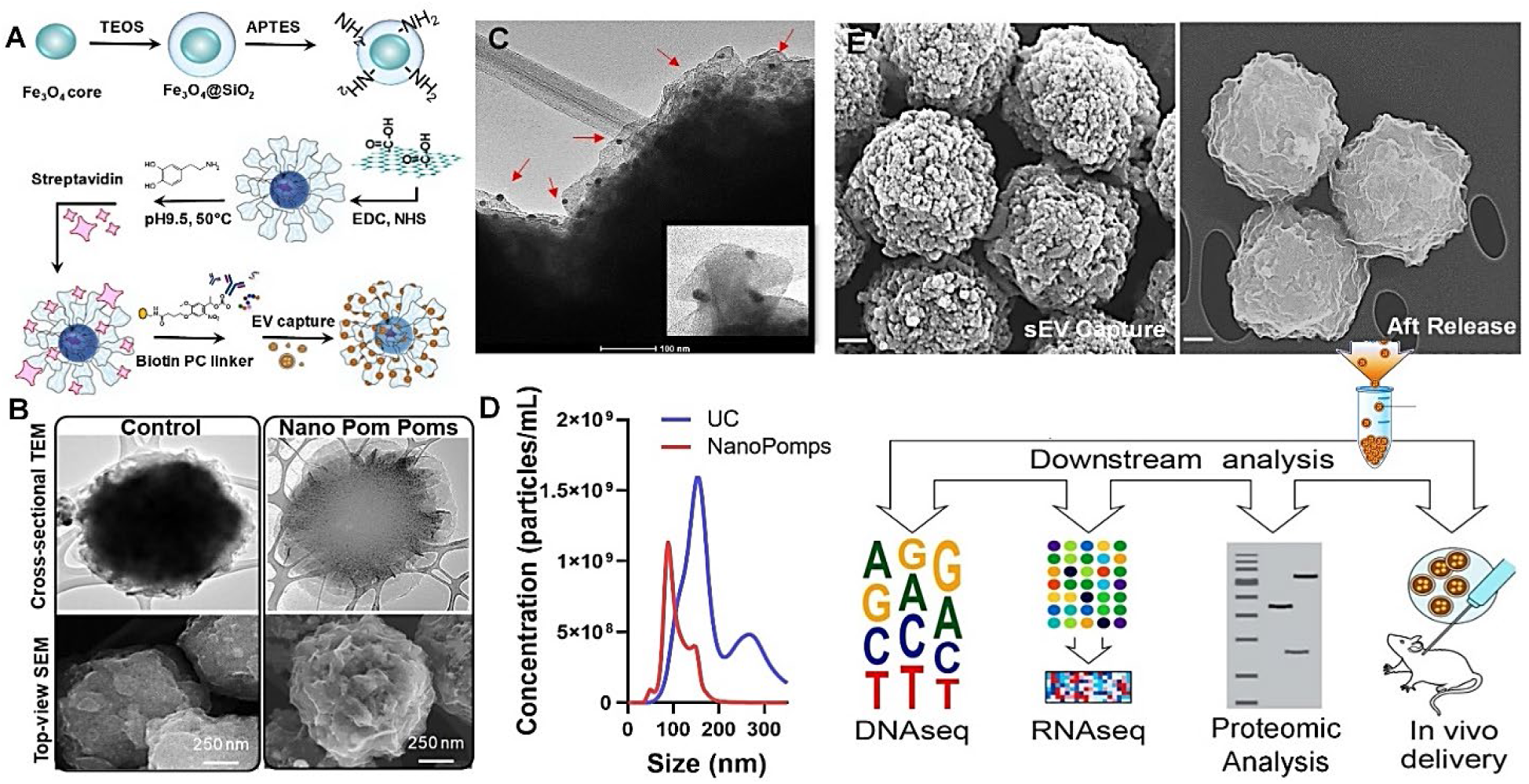
Nano pom-poms fabrication for highly specific sEVs isolation and multi-omic biomarker analysis. (**A**) Schematic illustration of the fabrication of Nano pom poms. (**B**) TEM and SEM images showing the unique 3D nano-scale flower pom-poms morphology compared to commercial immunomagnetic beads. (**C**) TEM imaging of captured sEVs fully covering Nano pom-poms surface. The captured EVs are confirmed by antiCD63 gold nanoparticle immune TEM imaging. The insert shows the captured single EV in the size range of ∼100 nm with three gold nanoparticles bound (∼10 nm). (**D**) Nanoparticle tracking analysis of NanoPoms isolated sEVs with much narrower size distribution in comparison with UC isolated EVs. (**E**) SEM images showing the dense sEVs are captured on Nano pom-poms, and can be completely released via on-demand photo-cleavage. After release, intact sEVs can be harvested for downstream multi-omic analysis including next generation sequencing of DNAs, RNAs, western blotting and proteomic analysis, as well as the in vivo study.

### NGS analysis of somatic DNA mutations carried by urinary Evs

Detecting DNA mutations carried by urinary tumor EVs is emerging, yet challenging, due to the needs of highly pure sample preparation for sensitive detection. We analyzed the bladder cancer (BC) patient urine samples prepared by both NanoPoms, UC, and commercial bead approaches for isolating urinary EVs, with control group from healthy individuals. The NGS GeneRead AIT panel was used to identify the most cancer relevant 1,411 variants. UC preparation was found insensitive to cancer relevant variant detection, as it requires much larger urine sample input (4 mL) with more than 100 ng EV DNAs to give detectable variant signals (Fig. 2a). We suspect that UC isolated EV DNAs contain more genes which are not specific to cancers. The PDGFRA variant (c.1432T>C, p.Ser478Pro) with 56.8% frequency was detected from a healthy individual in the control group using UC preparation, but not from NanoPoms preparation. In the BC disease group, NanoPoms prepared sEV enabled much enhanced detection sensitivity and specificity to BC relevant mutations including *KRAS, PIK3CA*, and *ERBB2*, which only consumed 1 mL urine sample with using about 10-50 ng sEV DNAs. However, commercial bead isolated EVs using the same urine sample input did not yield sufficiently enriched DNAs for sensitive detection of cancer relevant variants (Fig. 2a). In order to validate whether the gene mutations found in urinary EVs are from the urological tumor, we evaluated the matched patient tumor tissue. The NGS GeneRead analysis of tumor tissue cells showed the consistent mutations of *KRAS* and *ERBB2* which also were presented in the urinary EVs from the same BC patient. Although as one might expect, more mutations were detected in the tumor tissue, including *MTOR* and *BRCA1*; however, the pathogenic *PDGFRA* variant (c.1939A>G, p.Ile647Val) was found in the urinary EVs from both UC and NanoPoms preparations, but not in the tumor tissue cells (Fig. 2b). It is worth mentioning that the *PDGFRA* variant (c.1939A>G, p.Ile647Val) has been recognized as the tumor marker from the bladder urothelial carcinoma and the gastrointestinal stromal tumor(*31, 32*), which indicates that urinary sEVs could be a good biomarker resource and surrogate for tumor cells.

**Fig 2.**
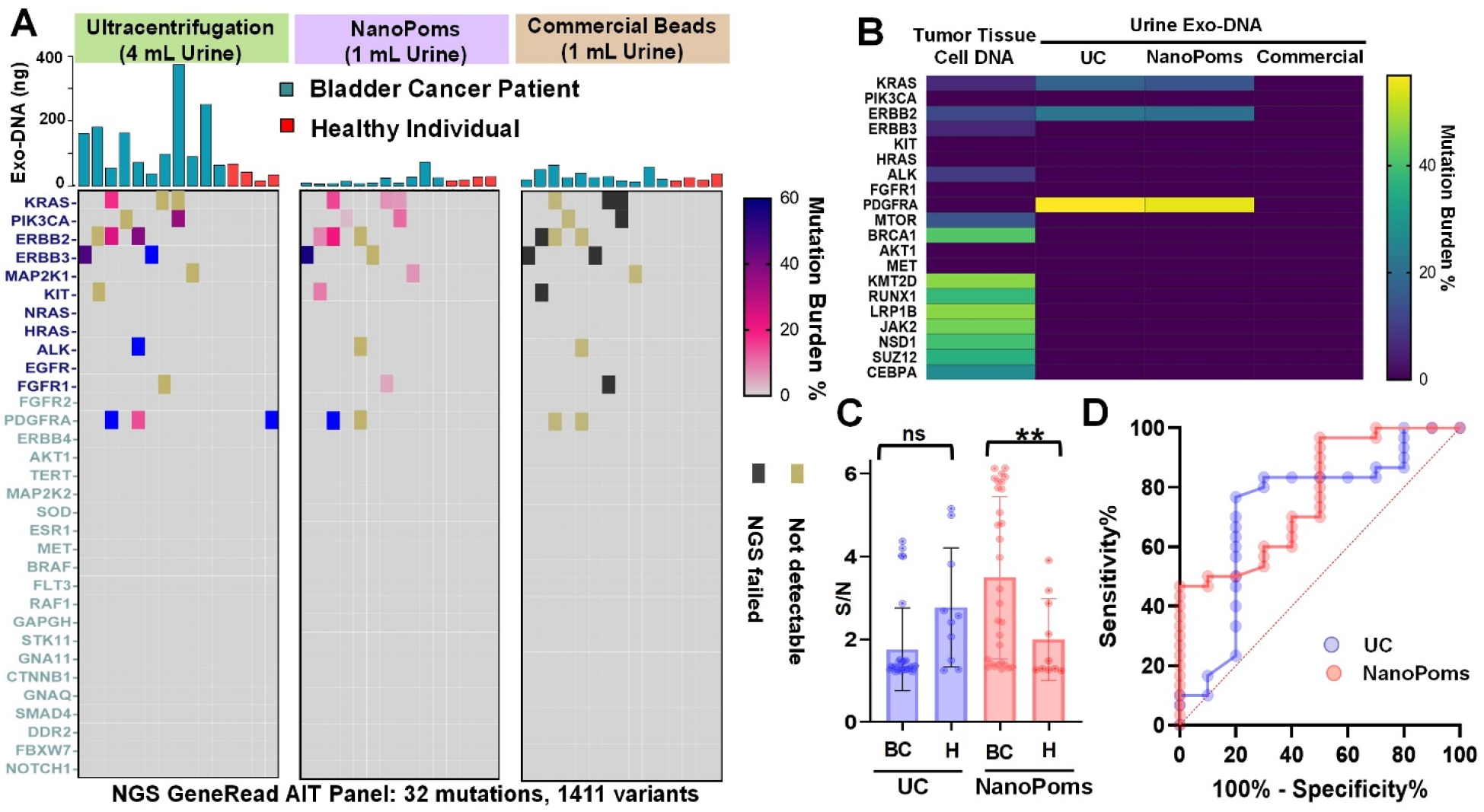
The NGS analysis of somatic DNA mutations from bladder cancer patient urinary EVs. **(A)** The DNA NGS analysis of 11 BC patient urine samples with 4 healthy individuals as the control group using GeneRead AIT panel. The EVs were prepared in parallel by UC, NanoPoms, and commercial bead approaches to extract total DNAs shown in the bar graphs. The most frequent 1,411 cancer relevant variants were sequenced. **(B)** The NGS GeneRead analysis of tumor cell DNAs from the matched BC patient tumor tissue, compared with urinary EV DNAs prepared by UC, NanoPoms, and commercial beads. **(C)** The droplet digital PCR analysis of *EGFR* (Thr790Met) extracted from purified EVs using both NanoPoms (pink dots) and UC (blue dots) approaches from 30 bladder cancer patient urine samples with 10 healthy individuals as the control group. **(D)** Receiver operating characteristic (ROC) analysis of ddPCR detection of *EGFR* showing significant diagnostic performance from NanoPoms approach compared with UC preparation. The a.u.c (Area Under the Curve) for NanoPoms preparation is 0. 78 with *p* =0.01. The a.u.c. for UC preparation is 0.71 with *p* =0.04.

We also analyzed urinary EV-derived DNA mutations using droplet digital PCR (ddPCR) from both UC and NanoPoms preparations. A total of 30 bladder cancer patient urines were analyzed with 10 healthy individuals as the control group. With the same EV DNA input (10 μg), *EGFR* (Thr790Met) and *TERT* (C228T and C250T) were detected. We observed much higher signal amplitudes from NanoPoms prepared EV DNAs than that from UC approach (Fig. 2c). The average patients’ EGFR Wt copy number is 3185.4 ± 468.3 from NanoPoms approach, which is 12.8-fold higher than that from UC approach (248.9 ± 46.4) with 3-fold higher mutation detection efficiency (Fig. S4). The overall detection signal to base ratio from patient group is statistically higher than that from control group (Fig. 2c), indicating the significant diagnostic value (Fig. 2d) for developing liquid biopsy and non-invasive diagnosis of BC using urinary sEVs from NanoPoms preparation. In contrast, UC-based EV preparation is unable to differentiate patient group from the healthy control group (*p* >0.05, Fig. 2c).

Interestingly, we also observed *EGFR* heterozygous mutations in three BC patients while conducting ddPCR analysis of NanoPoms prepared urinary sEV DNAs (Fig. s5). In contrast, UC isolates from the same patients 2 and 3 did not show such heterozygous mutation (Fig. 3a and Fig. s5). In order to further validate this observation, we obtained the matched patient plasma and buffy coat with white blood cells (WBC) as the control. NanoPoms preparation allows to pull out marker specific sEV populations based on the exosomal surface markers (CD9, CD63, and CD81) to match urinary EV populations, which avoids the interferences from other microvesicles or non-disease associated vesicles. Afterwards we used Sanger sequence to confirm the presence of the *EGFR* heterozygosity for three patients. Results were consistent with ddPCR analysis from NanoPoms preparation. As expected, the *EGFR* heterozygosity was not detected from matched patient WBCs wide-type control. These results clearly support that marker specific capture and release enabled by NanoPoms method can significantly enrich tumor-associated sEVs for sensitive mutation detection. Although the UC preparation yields larger numbers of vesicle particles, their specificity and purity to tumor-associated sEVs are much less than NanoPoms preparation.

**Fig 3.**
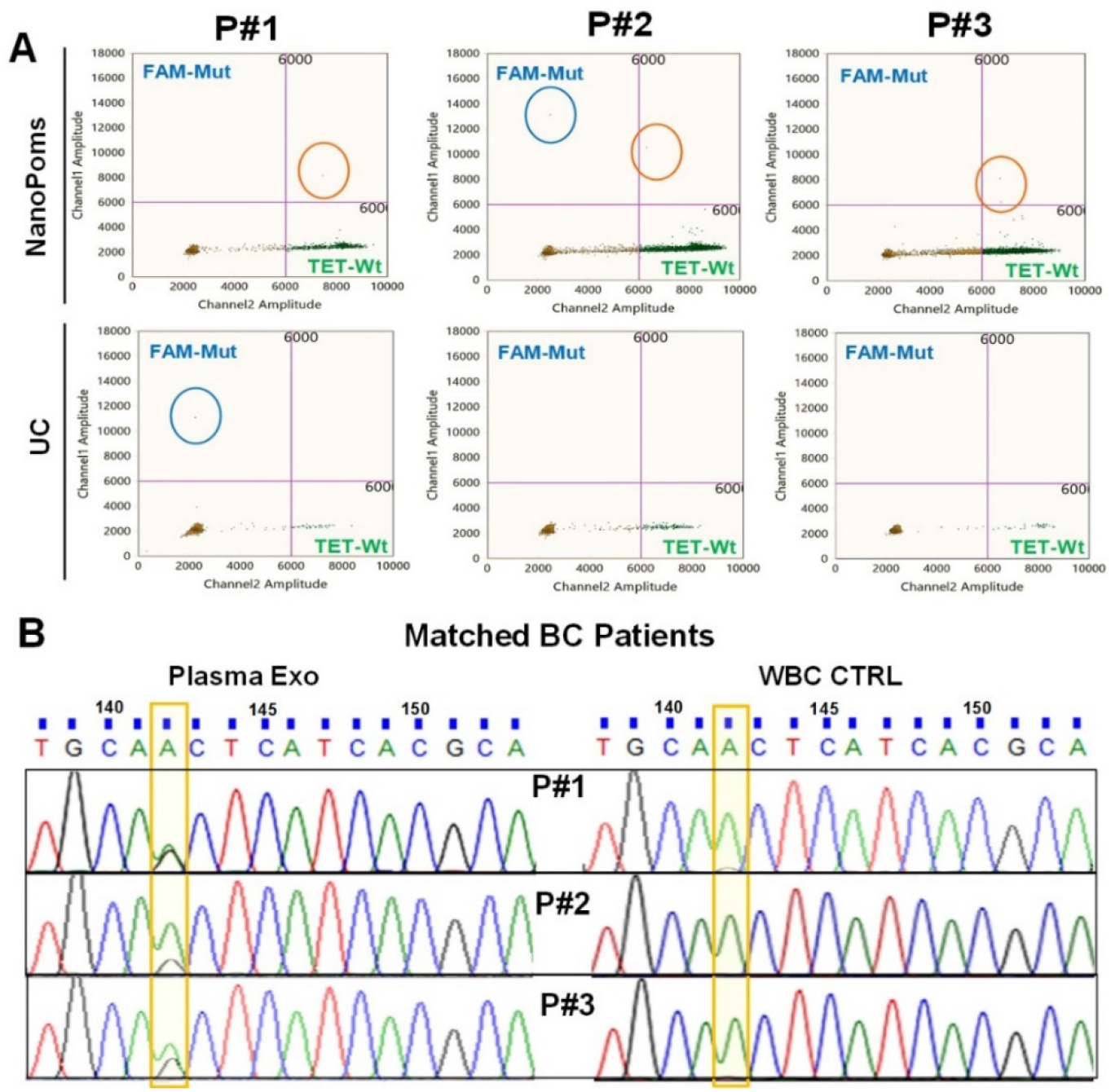
NanoPoms prepared sEVs enable highly sensitive detection of heterozygosity using ddPCR. **(A)** The ddPCR detection of *EGFR* (Thr790Met) from NanoPoms prepared urinary sEVs in three BC patients, compared with UC preparation. **(B)** Sanger sequence validation of *EGFR* heterozygosity from NanoPoms prepared plasma sEV in matched patients in Fig. 3A. Genes from the corresponding patients’ white blood cells (WBC) are the wild-type control.

### NGS analysis of urinary EV small RNAs

Analyzing RNAs within urinary EVs has been emerging with needs for non-invasive, early detection, and timely medical checkup of BC(*33, 34*). EV long non-coding RNAs (lncRNAs) *PVT-1, ANRIL* and *PCAT-1* have been reported as the novel biomarker in BC diagnosis(*35-38*). However, NGS profiling of microRNA from tumor derived urinary EVs has not been exploited. In this study, we analyzed urinary EV microRNA NGS profiles from both BC and healthy individuals. The distribution of EV small RNA categories from NanoPoms preparation showed more lncRNAs in both the BC group and healthy control group (42% from NanoPoms vs. 18.9% from UC) (Fig. 4a, Table s2). In contrast, UC preparation leads to the higher percentage of tRNA. Although the exact role of EV lncRNAs is not well understood yet, several studies have showed exosomal lncRNAs are novel biomarkers in cancer diagnosis and are highly associated with cancer progression and cellular functions(*39-41*). Currently, only a small number of lncRNAs has been investigated which partially due to the variation and uncertainty imposed by EV preparation methods(*42*). We further look into the top 100 miRNAs expression profiles as shown in Fig. 4b. The heatmap clustering analysis indicates the clear differentiation between BC group and healthy control from NanoPoms sEV preparation, in contrast to UC preparation. We also investigated the influence of photo cleavage process on the integrity of overall EV miRNAs (Fig s6), and did not observe significant differences in heatmap profiles with and without photo cleavage process. The NanoPoms sEV preparation leads to the significant differentiation ability between BC and healthy control groups, in contrast to UC approach.

**Fig 4.**
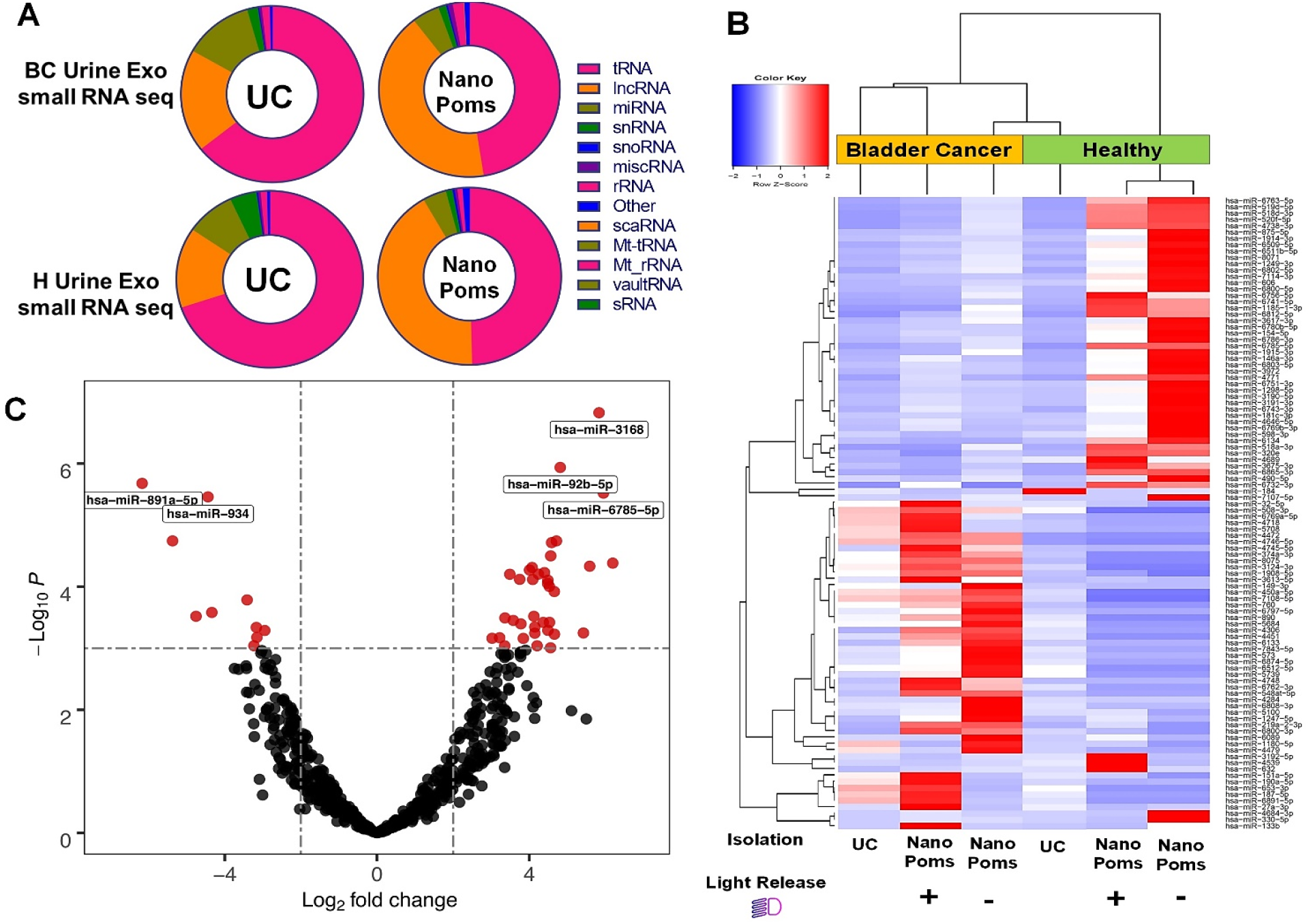
The NGS analysis of small RNAs from bladder cancer patient urinary EVs. **(A)** The distribution of small RNA categories from both NanoPoms and UC prepared urinary EVs in BC patients and healthy individuals. **(B)** Heatmap with dendrogram clustering analysis depicts the top 100 highly expressed miRNAs from urinary EVs isolated from both BC patient and healthy individual using UC, NanoPoms, and NanoPoms without light release process. Red color indicates a higher expression z-score. Hierarchical clustering was performed, using the Spearman correlation method. NanoPoms isolation approach with or without light release processes have been clustered together due to higher similarities in their transcript expressions. **(C)** Volcano plot analysis depicts the most biologically significant urinary EV miRANs with large fold changes identified by using NanoPoms preparation compared to UC preparation. Top 5 highly significant miRNAs are labeled in plot, which are from NanoPoms preparation.

In order to further interpret urinary EV miRNA profiles and characterize the influences imposed by EV sample preparation steps, we used the volcano plot to analyze the statistical significance (*P* value) versus fold-gene expression changes from both UC and NanoPoms EV preparations. It is interesting to note that top 10 miRNAs were highly enriched from the NanoPoms preparation, including hsa-miR-3168, hsa-miR-92b-5p, hsa-miR-891a-5p, hsa-miR-934, and hsa-miR-6785-5p (Fig. 4c and Table s3). We searched the reported miRNA functions and found those miRNAs were reported as the cancer relevant markers specifically sorted into exosomes (Table s3). For instance, hsa-miR-3168 has been reported to be enriched in exosomes via a KRAS-dependent sorting mechanism in colorectal cancer cell lines(*43*) and is known as the melanoma mature miRNA(*44*). The miR-92b-5p has been found to play a critical role in promoting EMT in bladder cancer migration(*45*). The hsa-miR-934 is an essential exosomal oncogene for promoting cancer metastasis(*46*). NanoPoms sEV preparation offered much higher molecular relevance for identifying tumor associated biomarkers, which is crucial for exploring more specific targetable cancer biomarkers.

### Proteomic analysis of urinary EV proteins

The urinary protein biomarkers could enable highly significant clinical values for the cystoscopic evaluations in BC diagnosis. EDIL-3 (Epidermal growth factor (EGF)-like repeat and discoidin I-like domain-containing protein 3) and mucin 4 (MUC 4) both have been reported in EVs purified from BC patient urines(*47, 48*). We selected generic EV markers CD9, CD63, and TSG101, as well as the EDIL-3 and MUC4 for Western blotting analysis of urinary EV proteins prepared by UC and NanoPoms approaches, with the human bladder carcinoma cell line HTB9 as the control (Fig. 5a). The generic EV markers CD9, CD63, and TSG101 were consistently expressed in urinary EVs, HTB9 cells and their EVs, which indicates consistent isolation of EVs. There is no significant difference between two preparation methods. The expression level of EDIL-3 is significantly higher in BC patients than healthy individuals, but not in the tumor cell line or their EVs from conditioned media. MUC4 protein marker was only observed in the human urinary EVs and HTB9 EVs, but not in HTB9 cells. This observation supports the previous report that EDIL-3 and MUC 4 are highly promising biomarkers in developing urinary EV-based BC diagnosis and prognosis tests(*47*).

**Fig 5.**
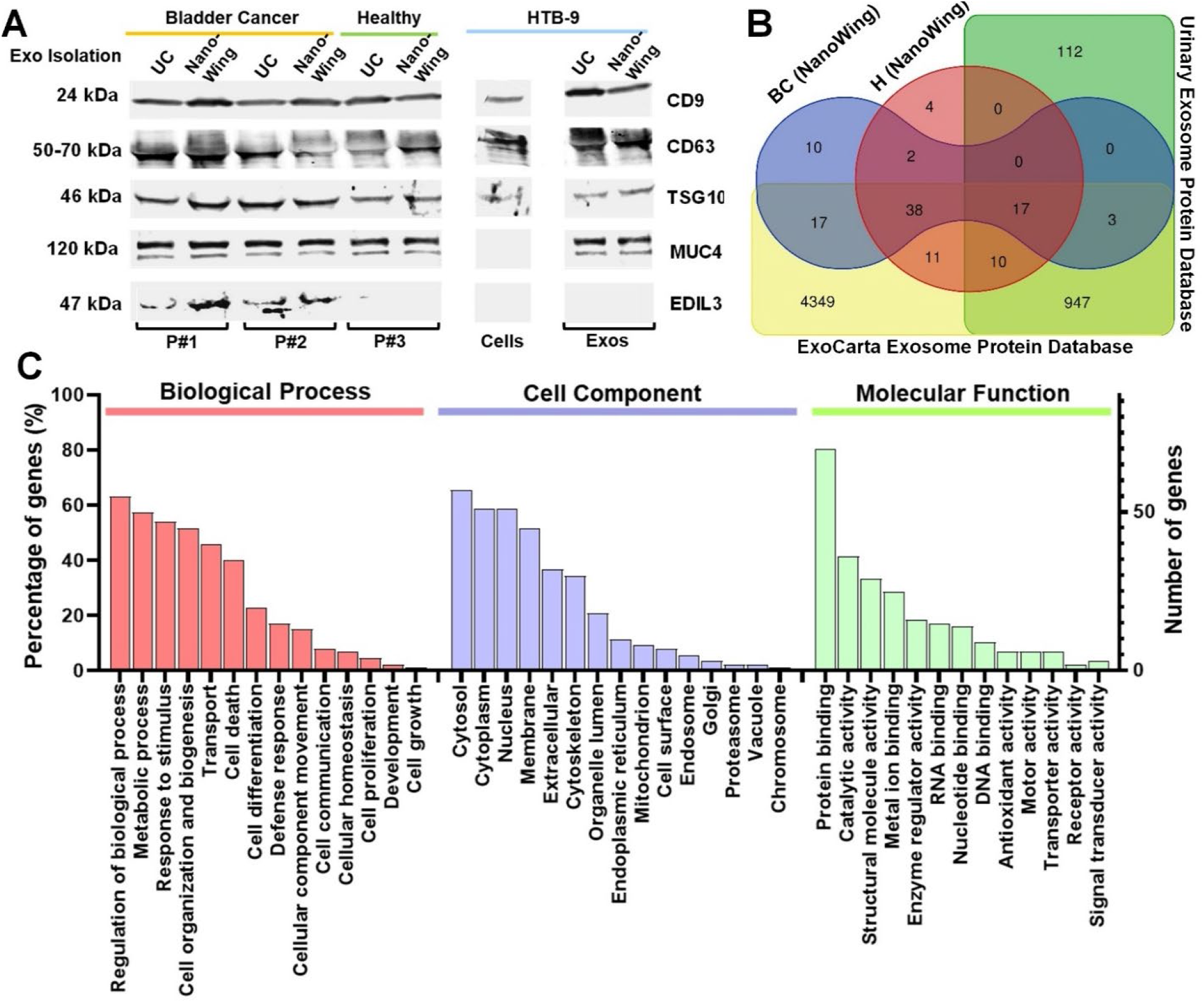
The proteomic analysis of bladder cancer patient urinary sEVs. **(A)** Western blotting analysis of urinary EV proteins prepared by both NanoPoms and UC approaches. Two BC patients and one healthy individual urine samples were used with HTB9 cells and their EVs from conditioned media as the control. Protein loading amount is applied consistently between samples (∼5 μg). **(B)** Venn diagram illustrates the relationship of proteomes from BC and healthy urinary sEVs prepared by the NanoPoms approach, with references from ExoCarta Exosome Protein Database and the Urinary Exosome Protein Database. **(C)** Gene Ontology enrichment analysis of differently expressed proteins from NanoPoms prepared BC urinary sEVs. Most abundant items are listed in biological process, cell component and molecular function, respectively.

The proteomic profiling of urinary EVs from NanoPoms preparations was shown in Fig. 5b, and identified proteins were compared with the ExoCarta Exosome Protein Database and the Urinary Exosome Protein Database. Several proteins associated with exosome biosynthesis were observed, such as proteins PIGQ and PAPD7 involved in Golgi apparatus, the cytosol protein S100-A7 and A9 found within the exosome lumen which is engaged with natural membrane budding process during multivesicular body formation. We also observed a diverse group of cytosolic enzymes (glyceraldehyde-3-phosphate dehydrogenase) and cytoskeletal constituents (actin, Beta-actin-like protein 2 ACTBL2, and myosin-9). Although the majority of proteins are shared identifications within BC patient and healthy control groups (∼65%), as well as the databases we used, interestingly, we found 10 proteins which are uniquely identified only from BC patient using NanoPoms preparation (Table s4). Those proteins have previously been reported to be associated with bladder cancer metastases, including *IRAK4(49), KRT23(50)*, and *RALGAPA2(51)* (full list in Table s4). Also 4 proteins were found uniquely in the healthy group using NanoPoms preparation, but not reported by ExoCarta and Urinary Exosome Protein Databases. From the Human Protein Atlas database (https://www.proteinatlas.org/), those proteins are intracellular and associated with vesicles, Golgi apparatus, and secreted pathway. The identifications are broadly consistent with that expected for exosomes and compatible with other researchers’ investigations(*52*). Approximately 35% of proteins do not overlapped between the BC patient and the healthy control, which further support the utility of NanoPoms prepared sEVs for aiding the diagnosis of BC.

Identified proteins were classified by encoding genes which indicate the majority are located within membranous vesicles, cytosol, cytoplasm, and the cytoskeleton, and some are located in Golgi (Fig. 5c). The biological processes associated proteome revealed significant associations with regulation of biological process, metabolic process, response to stimulus, cell organization and biogenesis, transport, and the cell death. The protein binding molecular function from this proteome is dominant. Results exhibit good specificity to exosomal proteome, indicating NanoPoms preparation could provide a pure and high-quality exosome type sEVs, which facilitates the important research area in EV proteomics and multi-omics.

### In vivo biodistribution study of NanoPoms prepared sEVs

The NanoPoms preparation of sEVs via marker specific capture and release is able to collect intact, pure and homogenous sEV subtypes. Due to the on-demand, light-triggered release process, the molecular engineering, such as the surface modification, drug loading, or dye labeling, can be implemented to immunomagnetically captured EVs before washing and releasing. This protocol avoids the redundant post purification of small molecules from isolated sEVs, which is often challenging and causes contaminations. For instance, the remaining free dye during in vivo tracking of EVs could cause false signals with longer distribution half time, unspecific staining or tissue accumulation(*53*). In this study, we prepared sEVs from bladder tumor HTB9 cells and non-malignant HEK cells with DiR labeling for intravenous tail injection into BALB/cJ mice. The buffer solution from beads washing step (without EVs) was used as the negative control. From these representative images in 24, 48, and 72 hr time intervals post injection, the fluorescent whole mouse imaging is not be able to provide enough precision to describe the levels of sEV distributions in tissue organs (Fig 6a). Thus, the organs were harvested and imaged ex vivo in the time intervals of 48 and 72 hr to minimize signal interference (Fig. 6 b and c). To rule out of the signal originating from the blood in the organs or from the free dye, we normalized the EV tracking signal with the negative control signal to affirm the in vivo EV tacking. In fact, the negative control images did not show much detectable signals indicating no remaining free dye background signal during in vivo tracking of sEVs. By further observing the harvested organs, HTB9-derived sEVs exhibit different biodistribution profile in lung, liver, kidney, spleen, heart, and brain, as compared to sEVs isolated from the non-malignant HEK293 cells. sEVs prepared from the HTB9 tumor cells were more concentrated in the liver and spleen with gradually increased intensity from 48 hrs to 72 hrs post injection. In contrast, non-malignant HEK293-derived sEVs tend to spread from liver to lung and spleen after 48 hrs post injection. Although HTB9 sEV biodistribution profile has not been reported elsewhere previously, the HEK293 sEV biodistribution profile is consistent with reported study in C57BL/6 mice(*53*). Fig. 6c provides the repetitive and quantitative analysis of biodistribution pattern over time. The results potentially indicate the distinctive biodistribution profile from cancer-associated sEVs which could be very important for understanding tumor cell-mediated communications within the microenvironment. Currently, substantial efforts have been made for using sEVs as therapeutic agents or delivery vehicle in vivo. Thus, being able to reproducibly prepare pure and homogenous sEVs is critical for maintaining consistent biodistribution patterns.

**Fig 6.**
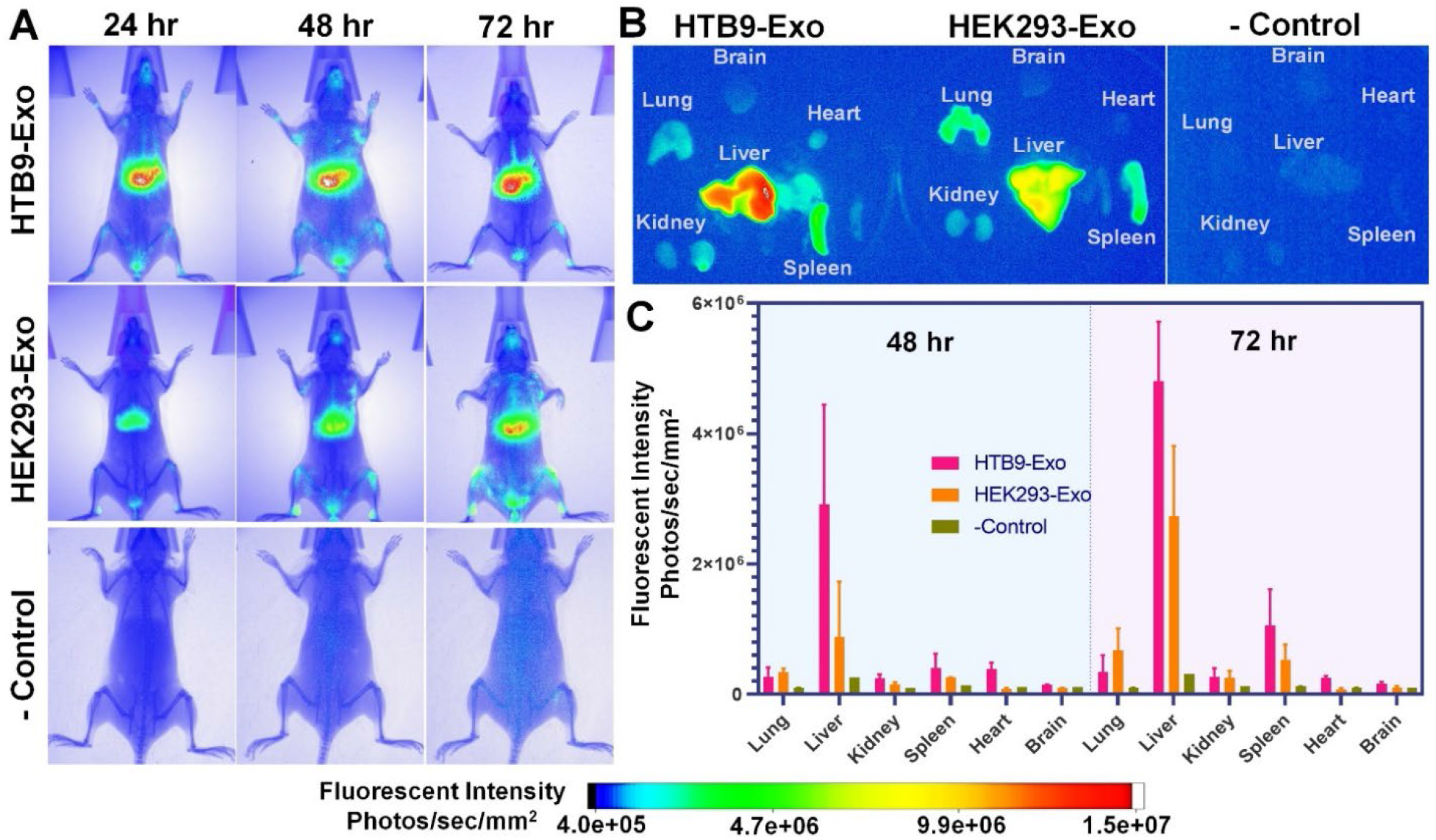
In vivo biodistribution analysis of NanoPoms prepared sEVs. **(A)** Representative IVIS images at 24 hours, 48 hours, and 72 hours post-injection of live mice. The HTB9 tumor cell derived sEVs and non-malignant HEK cell derived sEVs with DiR labeling (2.0 ×10^9^ particles/ml) were prepared by NanoPoms approach for intravenous tail injection into BALB/cJ mice. The buffer solution without EVs was used as the negative control. **(B)** Representative IVIS images of harvested organs (lung, liver, kidney, spleen, heart, and brain) at 48 hours and 72 hours post injection from mice. **(C)** The fluorescence signals normalized with negative control from IVIS images in each organ harvested at 48 hours and 72 hours post injection (n=2, mean ± SD).

## Discussion

All living cells secret extracellular vesicles (EVs) which are diverse populations with heterogeneous molecular functions(*54-56*). Recent and substantial researches have shown the heterogeneity of EVs(*17, 57-61*) in terms of density, molecular cargos, and morphology, which are even released by a single cell type(*62-64*). Our recent study also observed that molecular packaging of secreted EVs or exosomes is highly variable upon the change of cellular culture environment as well as surrounding community(*65*). Thus, the more advanced analytical methods are urgently needed to be able to decipher such heterogeneity in precision. Additionally, for therapeutic delivery, the well-defined molecular components from the homogenous EV population is also critical to precisely maintain controllable biodistribution pattern and delivery behavior(*66*). Due to the unique 3D nano pom poms structure and specific marker defined capture-release process, our developed isolation approach can prepare pure and homogenous sEV subpopulations which enrich tumor associated biomarkers.

The NGS analysis in our study demonstrated that DNAs isolated from NanoPoms prepared sEVs are enriched for tumor-associated DNA mutations which are highly relevant to the bladder cancer and comparable to matched patient tumor tissue cells (Fig. 2). The ddPCR analysis also confirmed such performance with significantly higher detectable copy numbers (Fig. 2c and Fig. S4 and s5). The heterozygosity also can be readily detected from very low level of copy numbers in NanoPoms prepared sEV DNAs, as confirmed by Sanger sequencing with matched patient plasma and buffy coat (Fig. 3 a and b, Fig. s5). The NGS analysis of sEV RNAs prepared by NanoPoms is able to reveal the distinctive profiles between BC patient and healthy individual, in terms of RNA types and miRNA levels, in contrast to UC preparation (Fig. 4 a and b, Table s2). Most importantly, the top 10 miRNAs identified from NanoPoms sEV preparation are highly relevant as the important cancer markers specifically sorted into exosomes (Fig. 4 c and Table s3). This evidence further supports that specific cancer-associated biomarker are enriched in exosome type urinary sEV and can serve as surrogates for tumor cells.

The miRNAs represent the most dynamic nucleic acid cargos in EVs, which is relatively sensitive to external stimulus and changes. Thus, in order to gauge the impact of light release process on sEV isolation via NanoPoms approach, we compared miRNA profiles with or without light release process, which did not show statistically significant differences based on dendrogram clustering analysis (Fig. 4b and Fig. s6). The light release process also is able to ensure the specificity via releasing captured sEVs only to get rid of non-specific binders. This data supports the quality and integrity of NanoPoms prepared sEVs as a novel, rapid, and easy-to-use method. Currently, although urinary miRNA profiling is highly essential for BC diagnosis, such study and relevant database have not been fully established yet. NanoPoms based sEV sample preparation could potentially speed up this research direction by offering much simple and specific sEV preparation.

The urinary sEV cargos at the protein level from our study reveals the consistent expression of exosomal proteins CD9, CD63, and TSG101 from both patient urinary EVs and cell lines using UC and NanoPoms preparations. In contrast, EDIL-3 levels have been observed much higher in BC patient urinary EVs compared to healthy individuals which is consistent with reported literature(*47*), indicating the high quality preparation of exosome type sEVs using NanoPoms approach (Fig. 5a). Further, the proteomic profiling also supports that NanoPoms prepared urinary sEV proteins can be used to differentiate BC disease from healthy status (Fig. 5 b and c, and Table s4) with unique identification of pathogenesis relevant EV proteins, suggesting a promising avenue using NanoPoms prepared sEVs to develop non-invasive bladder cancer diagnosis.

In order to further prove the integrity and biological activity of NanoPoms prepared sEVs, in vivo biodistribution study exhibits distinctive distribution patterns between tumor-associated sEVs and non-malignant sEVs (Fig. 6). This result may indicate that different subtypes and sources of EVs could have impact on the performance of drug delivery while using EVs as the carrier. To date, the therapeutic potential of different subpopulations of EVs is not well known. It has been discussed that possibly only a small fraction of the EVs from a cell can mediate the therapeutic effects (*62*). Thus, the reproducible isolation of specific sEV subpopulations are essential to support the development of EV-based therapeutic delivery. However, current existing EV isolation strategies are still unable to reproducibly differentiate sEV subpopulations. Our NanoPoms approach for enriching exosome type sEVs with marker definition could open a new avenue for preparing pure and homogenous sEVs with improved diagnostic and therapeutic efficacy.

## Materials and Methods

### Fabrication and Characterization of Nano Pom Poms

The proprietary bead fabrication follows the protocol of Fe3O4/SiO2 core-shell-based particle method with surface anchored graphene oxide nanosheets via carboxamide covalent bonds and EDC/NHS chemistry, and further modified with (3-aminopropyl) triethoxysilane (APTES), polydopamine, and streptavidin (Vector Laboratories, SA-5000). Beads were washed with PBST then resuspend in 1 ml PBST and 0.09% NaN_3_ solution for storage at 4 °C. In this study we used the pan capture with a mixture of CD9, CD63, and CD81 antibodies for bead-conjugation. For in vivo biodistribution study, we used CD9 antibody conjugated NanoPoms to prepare HTB9 and HEK cells derived sEVs. After bead fabrication and conjugation, XPS analysis was used (PHI 5000 VERSA PROBE II) with an Al anode of the x-ray source (46.95 eV) and 100 µ X-ray beam size for operating at 23.2 W. The power of the source was reduced to minimize X-ray damage for analyzing EVs on bead surface.

The EV isolation from patients’ plasma, urine, or cow milk and conditioned cell culture media were performed by incubation of 100 ul antibody-beads complex with 1 mL of samples at 4 °C overnight. After washing, the photorelease was performed using Analytikjena UVP 2UV Transilluminator Plus at 365 nm wavelength at 4 °C for 15 min (∼6 mW/cm^2^). The UC isolation of EVs followed the well-documented protocols published previous(*65*). Briefly, to remove any possible apoptotic bodies and large cell debris, the supernatants were centrifuged at 10,000g for 30 mins, then transferred to ultracentrifuge tube (Thermo Scientific, USA) for ultracentrifugation at 100,000g for 70 min (Sorvall™ MTX150 Micro-Ultracentrifuge, USA), with second ultracentrifugation (100,000g for 70 min) for finally collecting EV pellets. The size characterization of EVs was performed using the nanoparticle tracking analysis (NTA) Nano-Sight LM10 (Malvern Panalytical). Post-acquisition parameters were adjusted to a screen gain of 10.0 and a detection threshold to 5. Standard 100 nm nanoparticles were used for calibration. Appropriate sample dilution in 1x PBS was evaluated before every measurement with five repeats for each measurement.

### sEV DNA extraction and NGS sequencing

Frozen urine samples were thawed overnight at 4 °C and pre-centrifuged at 4 °C 10,000g for 30 min to remove cell debris. By using NanoPoms isolation, the extracted sEVs were treated with DNase I before DNA extraction. The QIAamp DNA Mini Kit (Qiagen, 51304) was utilized to extract DNA from all EV samples. The addition of 1 μl of an aqueous solution containing 10 μg of carrier DNA (poly dA) to 200 μl Buffer AL was used to ensure binding conditions are optimal for low copy number DNA according to the manufacturer’s protocols. DNA was Eluted in 20 μL Buffer AE. DNA concentrations were measured using a Nanodrop platform at an absorbance at 260 and 280 nm subtracted by the background value of carrier ploy dA only.

The library preparation by targeted enrichment using Qiagen GeneRead QIAact AIT DNA UMI and GeneRead clonal Amp Q Kits, was subjected to next-generation sequencing (NGS) to generate FASTQ files (text-based format for storing nucleotide sequences). This test is a targeted NGS Panel that encompasses 30 genes and 1411 variants (*AKT1, ALK1, BRAF, CTNNB1, DDR2, EGFR, ERBB2, ERBB3, ERBB4, ESR1, FBXW7, FGFR1, FGFR2, FGFR3, FLT3, GNA11, GNAQ, HRAS, KIT, KRAS, MAP2K1, MAP2K2, MET, NOTCH1, NRAS, PDGFRA, PIK3CA, RAF1, SMAD4, STK11*) with variable full exon or partial region. The reads are mapped to the Homo_sapiens_sequence hg19 reference and variants identified using QIAGEN QCI-Analyze pipeline.

The extracted DNAs were amplified by PCR to detect the EGFR (P00533:p.Thr790Met) mutation. The sequences of primers for PCR were as follows: Primer F, 5’-ATGCGTCTTCACCTGGAA-3’; primer R, 5’-ATCCTGGCTCCTTATCTCC-3’. Primers were designed by Primer3Plus online. The PCR assay was performed with Promega GoTaq Flexi DNA Polymerase kit in a 50-μL mixture containing 10μL of 5× PCR buffer, 0.25μL GoTaq Flexi DNA Polymerase, 10 μM of each primer (IDT, USA) and 20 μL of DNA in an ABI PCR instrument (Applied Biosystems). The PCR conditions were as follows: Initial denaturation at 95°C for 2 min, followed by 35 cycles at 95°C for 15 sec, 54°C for 30 sec and 72°C for 40 sec, then a hold at 72°C for 5 min and a final permanent hold at 4°C. The 319 bp DNA size of PCR products were clarified by 1% agarose gel electrophoresis using 5 μL PCR products and remained DNA were purified by QIAquick PCR Purification Kit (Qiagen, 28104). The purified PCR products were sequenced by Sanger Sequencing approach (GeneWiz, USA) using the same primers above.

### sEV RNA extraction and NGS sequencing

The miRNeasy Mini Kit (Qiagen, 217004) was used to extract total RNA from all EV samples per manufacture’s protocols. The amount of 700 μL QIAzol lysis reagent was adapted according to the manual. To achieve a higher RNA yield, the first eluate of 30 μL was applied to the membrane a second time. Isolated RNAs were quantified by High Sensitivity RNA ScreenTape Assay using Agilent TapeStation 2200 (Agilent, 5067-5579, 5067-5580). Total RNA was stored at -80 °C until small RNA Library preparation. The QIAseq miRNA Library is prepared for Single Read 75bp sequencing, with UMI tag per manufacture’s protocols. After small RNA sequencing using Illumina MiSeq system, the Qiagen specific UMI analysis per the kit instruction was performed with details in supplementary information.

### Droplet digital PCR

A pair of probes and a pair of primers were designed to detect *EGFR* and *TERT* mutation respectively. Due to the short size of the probe, in order to increase the hybridization properties and melting temperature, Locked Nucleic Acid (LNA) bases were introduced on the bases indicated with a “+”. One probe was designed to recognize wildtype (5′-TET/T+CATC+A+C+GC/ZEN/A+GCTC/-3′ IABkFQ). The second probe was designed to recognize the *EGFR* (P00533:p.Thr790Met) mutation loci, (5′-6FAM/T+CATC+A+T+GC/ZEN/A+GC+TC/-3′ IABkFQ). Primers were designed to cover both side of detection loci. For TERT, a probe was designed to detect both C228T and C250T mutation as both mutations result in the same sequencing string(*67*), with (TERT Mut:/56-FAM/CCC+C+T+T+CCGG/3IABkFQ/). A second probe was designed to recognize the C228 loci, also containing LNA bases, (TERT WT, /5HEX/ CCCC+C+T+CCGG/3IABkFQ/). Probes and primers were custom synthesized by Integrated DNA Technologies (IDT). Amplifications were performed in a 20 μL reaction containing 1 × ddPCR Supermix for Probes (No dUTP),(Bio-Rad, 1863024), 250 nM of probes and 900 nM of primers and 8 μL EV DNA template. Droplets were generated using the QX200 AutoDG Droplet Digtal PCR System (Bio-Rad). Droplets were transferred to a 96-well plate for PCR amplification in the QX200 Droplet Reader. Amplifications were performed using the following cycling conditions: 1 cycle of 95°C for 10 minutes, then 40 cycles of 94 °C for 30 seconds and 60 °C for 1 minute, followed by 1 cycle of 98°C for 10 minutes for enzyme deactivation. Keep all ramp rate at 2°C/sec. QuantaSoft analysis software (Bio-Rad) was used to acquire and analyze data.

### Western blotting and Proteomic analysis

The 5 mL of each urine sample for two patients and one healthy control were used for EV isolation and subsequent Western blot analysis. 40 mL of HTB-9 conditional cell culture media and 40 mg cell pellets were also used as controls in this study. Samples were lysed in 1× RIPA buffer supplemented with protease inhibitors for 15 min on ice. Only cell sample were ultrasonicated for 1 min. Protein concentration was quantified using Micro BCA Protein Assay Kit (Thermo Fisher, 23235). The absorbances were read at 562 nm on a Synergy H1 reader (BioTek). All sample concentration were adjusted to 0.1 μg/μL. Western blotting was performed under reducing conditions (RIPA buffer, β-mercaptoethanol and Halt Protease Inhibitor Cocktail, EDTA-Free) at 95 °C for 5 min. 20 μL of protein lysate, each, were loaded onto 4-20% Mini-PROTEAN TGX Precast Protein Gels (BioRad, 4561093). The separated proteins were transferred to a PVDF membrane (BioRad, 1620218). After blocking the membrane in Intercept (PBS) Blocking Buffer (LI-COR, 927-70001) for one hour at room temperature, it was incubated over-night with the primary antibody at 4 °C, followed by another incubation with the secondary antibody for half hour at room temperature. The following primary antibodies were used, all diluted in blocking buffer (1:1000): anti-CD9 (Thermo Fisher, 10626D), anti-CD63 (Thermo Fisher, 10626D), anti-EDIL3 (Abcam, ab88667), anti-MUC4 (Abcam, ab60720), anti-TSG101 (Invitrogen, PA5-86445), anti-ANXA7 (LSBio, LS-C387129-100). The secondary anti-mouse and anti-rabbit IRDye 800CW antibodies (LI-COR, 926-32210 and 926-32211) were applied in 1:15,000 dilution. Imaging were performed by LI-COR Odyssey CLx system.

Urinary EV pellets resultant from ∼2 mL of urine from both bladder cancer patients and healthy individuals were reconstituted in 400 µL of M-PER Mammalian Protein Extraction Buffer (Thermo) supplemented with 1× Halt Protease Inhibitors (Thermo) and sonicated in an ultrasonic water bath for 15min. Lysates were exchanged into ∼40 µL of 100mM triethylammonium bicarbonate using Amicon Ultra-0.5, 3k columns (Millipore). Lysate were digested overnight with Trypsin Gold, Mass Spectrometry Grade (Promega) for subsequent HPLC-MS detailed in the supplementary material.

### SEM and TEM

sEV-particle complex was resuspended in 200 μL cold PBS solution. For electron microscope evaluation, EV-particle complexes were washed with pure water followed by the fixation in a 2% EMS-quality paraformaldehyde aqueous solution. 5 μL of sEV-particle mixtures were added to cleaned silicon chips and immobilized after drying EVs under a ventilation hood. Samples on silicon chips were mounted on a SEM stage by carbon paste. A coating of gold-palladium alloy was applied to improve SEM image background. SEM was performed under low beam energies (7 kV) on Hitachi SU8230 filed emission scanning electron microscope. For TEM, ∼5 μL of each sEV-particle complex was left to adhere onto formvar carbon coated copper Grid 200 mesh (Electron Microscopy Sciences) for 5 mins followed by 5 mins of negative staining with 2% aqueous uranyl acetate. Excess liquids were blotted by filter papers. Total grid preparation was performed at room temperature till totally air-dried under a ventilation hood for 25 mins. Images were acquired on the same day at 75 kV using Hitachi H-8100 transmission electron microscope.

### In vivo biodistribution analysis

The human bladder cancer cell line HTB-9 (ATCC, 5637) and the negative control of human embryonic kidney epithelial cell line HEK293(ATCC, CRL-1573) were cultured in DMEM and MEM respectively, supplemented with 10% normal FBS and 1% penicillin/streptomycin. Once the cell cultures reached ∼70% confluency, the media was replaced with fresh media containing 10% exosome-depleted FBS (Thermo Fisher, A2720803). The cells were cultured for an additional 72 h before the conditioned media were collected.

sEVs were isolated using NanoPoms approach. sEVs were incubated with 1 mM fluorescent lipophilic tracer DiR (1,1-dioctadecyl-3,3,3,3-tetramethylindotricarbocyanine iodide) (Invitrogen, D12731) at room temperature (RT) for 15 minutes. DiR-labelled sEVs or free DiR dyes were segregated using Amicon Ultra-15 Centrifugal Filter method. The 2.0 ×10^9^ particles/ml of isolated sEVs measured via NTA were used for each mouse injection. The 6-to 8-week-old female BALB/cJ mice were used. The animal IACUC protocols have been approved by the University of Kansas Institutional Animal Care and Use Committee with protocol number 258-01 and operated in the KU Animal Care Unit. Freshly purified DiR-labelled sEVs were injected through the tail vein for intravenous (i.v.) injection. The In-Vivo Systems (Bruker, USA) with high-sensitive CCD camera was used for collecting fluorescence, luminescence and X-ray images. Isoflurane sedated live mice were taken fluorescence and X-ray images prior to the animals were sacrificed, then main organs (brain, heart, lung, liver, kidney and spleen) were harvested for fluorescence imaging in 3 mins (excitation 730 nm, emission 790 nm), X-ray imaging (120 mm FOV, 1 min) and luminescence imaging (90 fov, 0.2 sec) at 24 h, 48h and 72 h time points, respectively. The data were analyzed using the Bruker MI software.

### Data statistics

All statistical tests were performed under the open-source statistics using GraphPad Prism software 8 (San Diego, CA), including heatmap, ROC analysis, and clustering analysis. The one-way ANOVA and t test were used. Differences are considered statistically significant at P < 0.05. *P < 0.05; **P < 0.01; ***P < 0.001. The Venn diagram was analyzed using open software from the Bioinformatics & Evolutionary Genomics. The bioinformatic analysis of small RNAs was detailed in supplementary material.

## SUPPLEMENTARY MATERIALS

Supplementary material for this article is available

## Acknowledgments

We thank Xinbao Hao for helping with the in vivo biodistribution imaging protocols. We also thank the assistant from Jennifer Hackett from the Genomic Core at the University of Kansas for library preparation, quality check and next-generation sequencing. We acknowledge the support of The University of Kansas Cancer Center’s Biospecimen Repository Core Facility staff, funded in part by the National Cancer Institute Cancer Center Support Grant P30 CA168524 (A.K.G.).

## Author contributions

Conceptualization: MH, YZ

Methodology: NH, ST, CZ, ZG, LX, ZP,AKG

Investigation: NH, ZG

Visualization: MH, ST, CZ

Supervision: MH, YZ, CZ, LX, AKG

Writing: MH, YZ, NH, AKG

## Data and materials availability

The authors declare that all data supporting the findings of this study are available within the paper and its Supplementary Information files. The DNA and RNA sequence data, as well as the protein proteomic data are made available from depositing to exosomal databases ExoCarta or upon request. Raw data regarding study is available for research purposes from the corresponding author on reasonable request.

### Supplementary Materials for

**Fig. S1.**
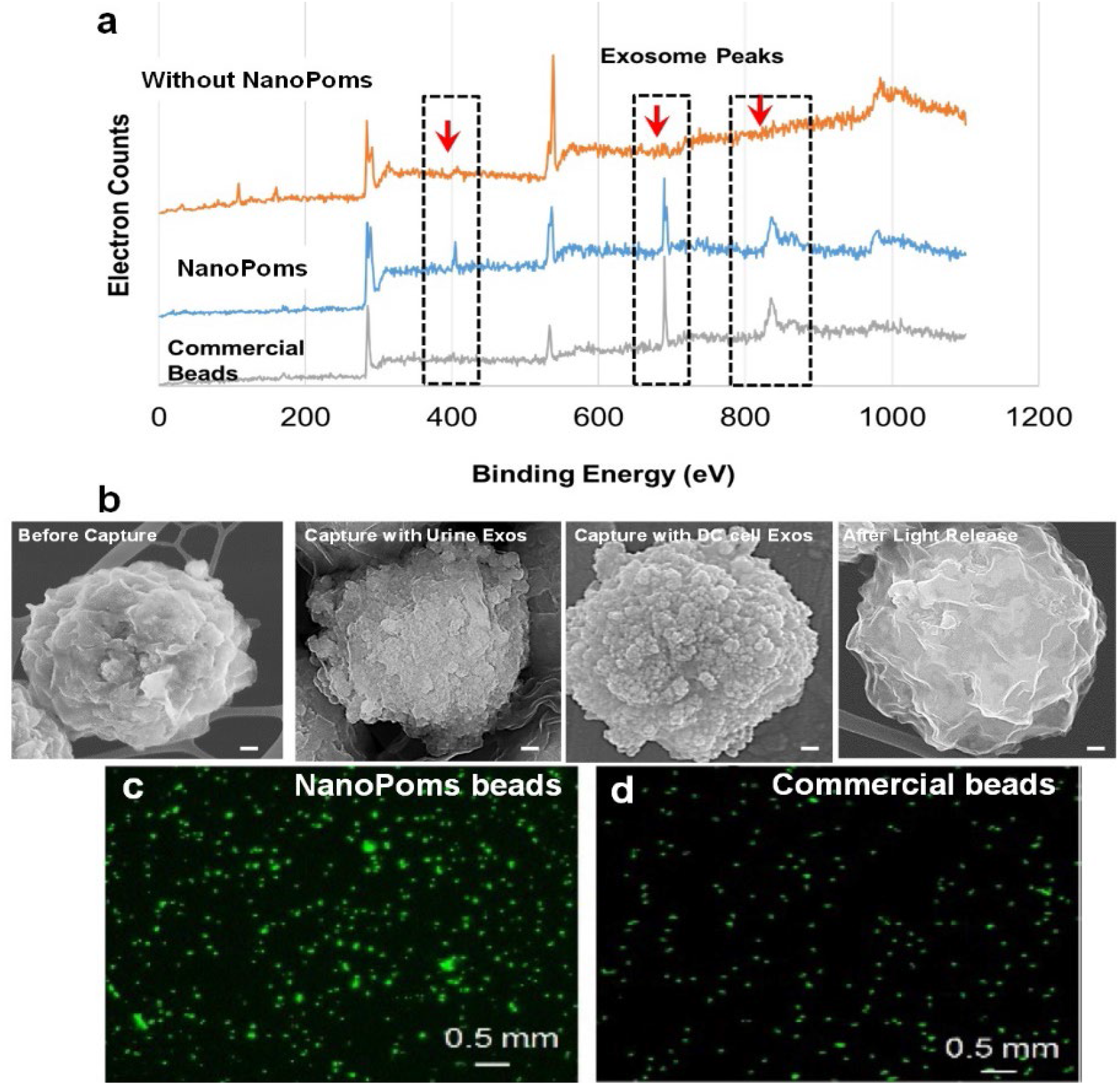
Characterization of Nano Pom Poms particles for specific capture and release of sEVs. **a)** X-Ray Photoelectron Spectroscopy (XPS) analysis of Nano pom-poms surface properties with extracellular vesicles captured. The bare magnetic particles without the 3D-structured nanographene sheet layers serves as the negative control. The commercial dynabeads were used as the positive control. **b)** The SEM images showing the surface morphology of Nano pom-poms before EV capture, after EV capture, and after release of captured EVs. The scale bar is 100 nm. The dense round small EVs were seen covering particle surface completely after capture. **c)** Fluorescence microscopic images showing the Nano pom-poms bound to FITC-biotin after conjugation with streptavidin, with dyna streptavidin beads as the positive control, which exhibits much brighter fluorescence from Nano pom-poms indicating more binding sites.

**Fig. S2.**
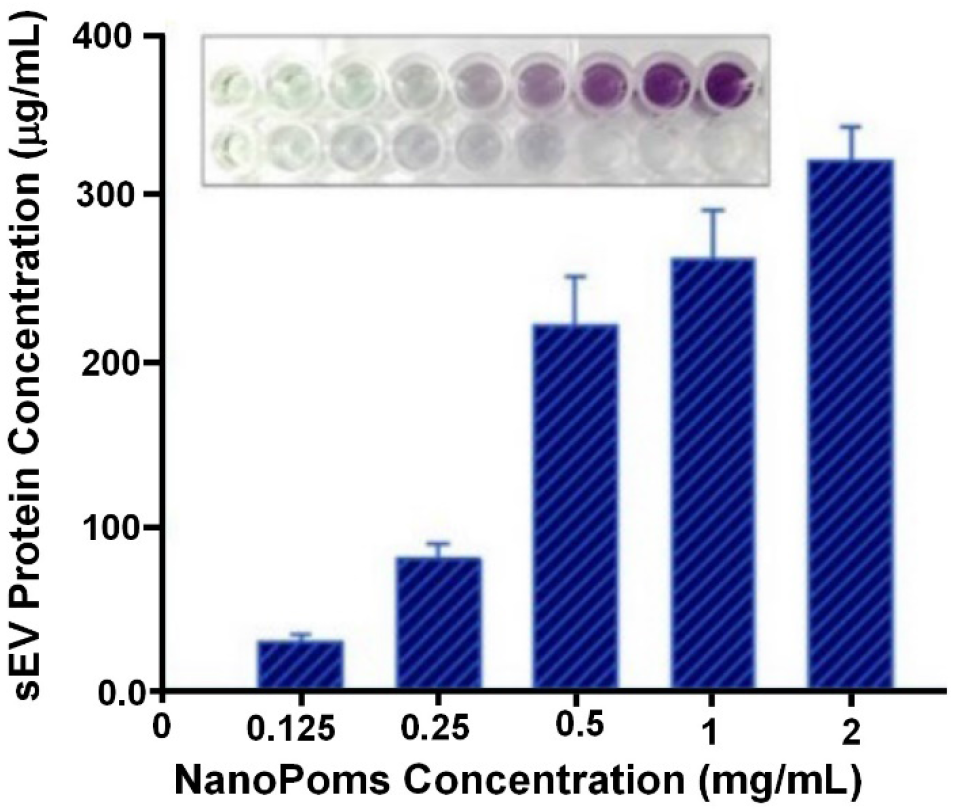
The optimization of Nano Pom-poms concentration used for isolating sEVs. The total protein concentration from isolated sEVs is measured by the Pierce BCA Protein Assay. The five repetitive measurements were performed for each data point with RSD < ∼5% (n=5). With increasing the amount of capture NanoPoms in 1 mL biofluids (here cow milk), more sEVs were isolated to gradually reach to the maximum. Once the available capture binding sites on the particle surface excess the number of overall EV particles, the increase of particle amount does not influence on the overall EV capture amount. Thus, the 1mg/mL of NanoPoms capture particles were used for appropriate capture efficiency ranging from ∼10^8^ to ∼10^13^ particles/mL.

**Fig. S3.**
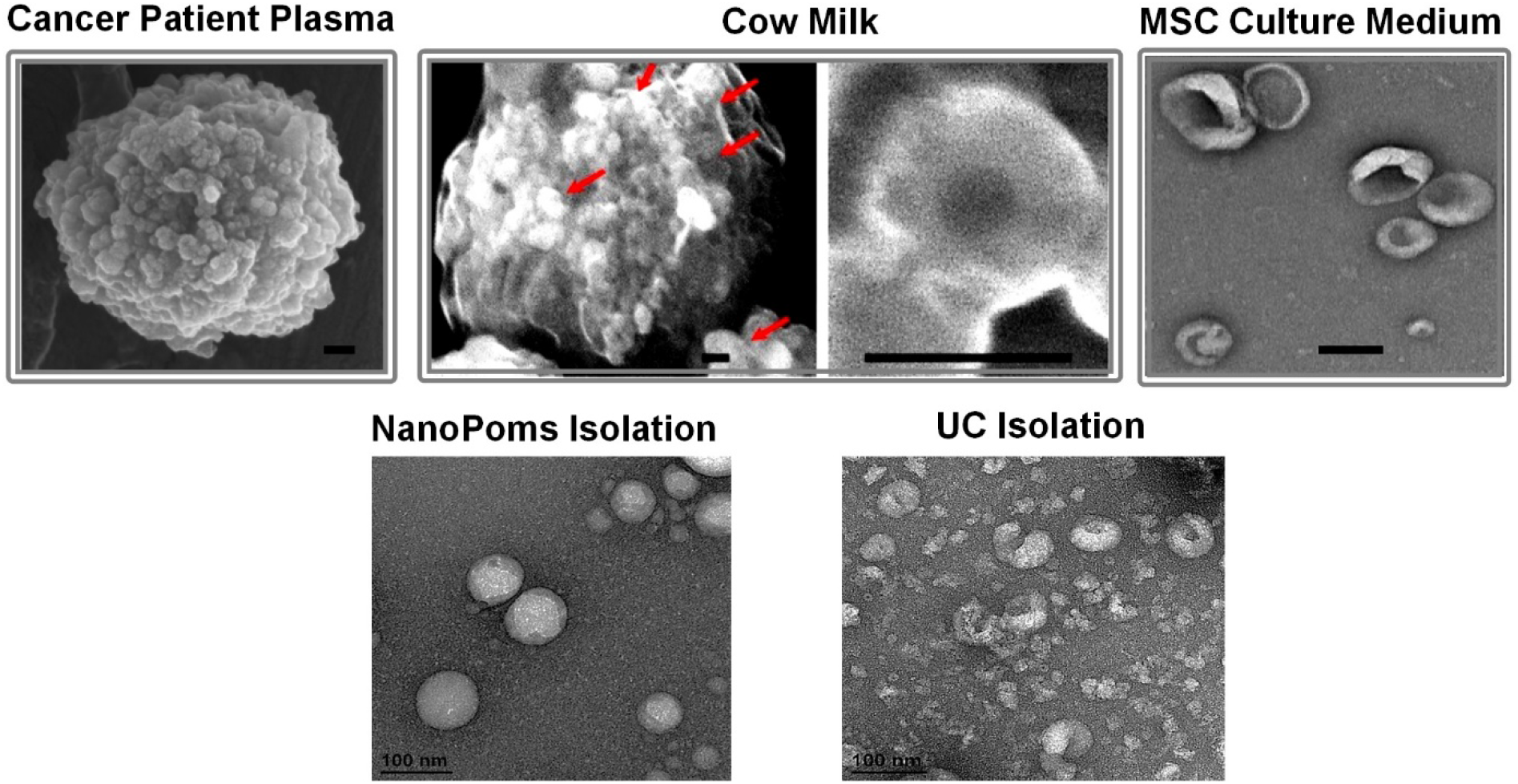
The SEM and TEM morphological characterization of sEVs isolated from a variety of biological fluids using Nano pom-poms. Top: The SEM images showing the morphology of Nano pom-poms captured sEVs from ovarian cancer patient plasma (left), cow’s milk with enlarged insert showing the classic cup shape (middle), released from the Wharton’s jelly mesenchymal stem cell culture medium (right). The scale bar indicates the 100 nm. Bottom: TEM images showing the much clean and uniform sEVs isolated from NanoPoms preparation from cell culture medium. In contrast, ultracentrifugation prepares EVs in a mixture with small aggregates and debris.

**Fig. S4.**
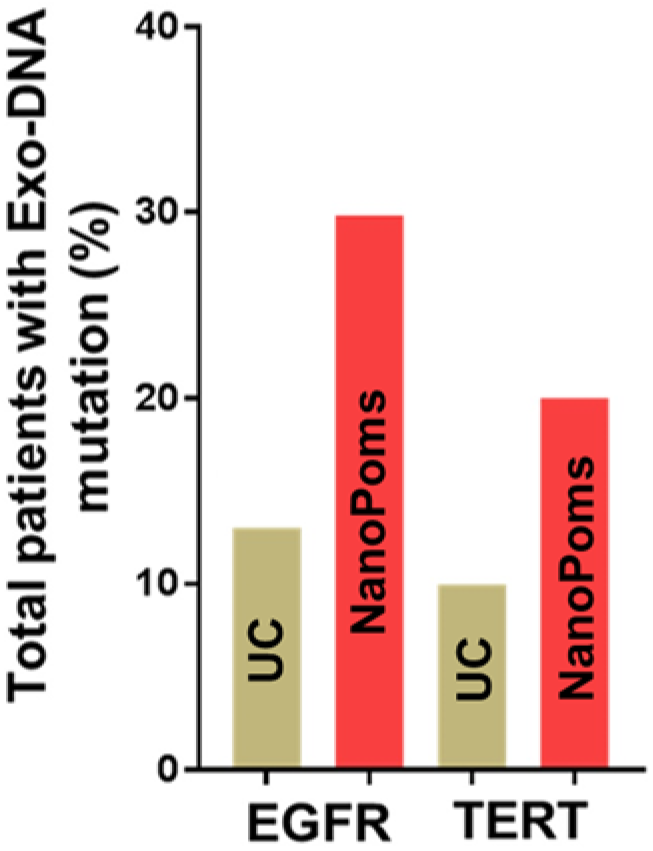
The droplet digital PCR analysis of NanoPoms prepared sEVs with substantially improved detection specificity and sensitivity. The ddPCR analysis of the mutation frequency *EGFR* (Thr790Met) and *TERT* (C228T and C250T) using DNAs extracted from either NanoPoms or UC prepared urinary EVs. The 30 bladder cancer patient urine samples were used. The same amount of EV DNAs (10 µg) were used as the sample input. The detection efficiency from NanoPoms was 3-fold higher as compared to UC approach.

**Fig. S5.**
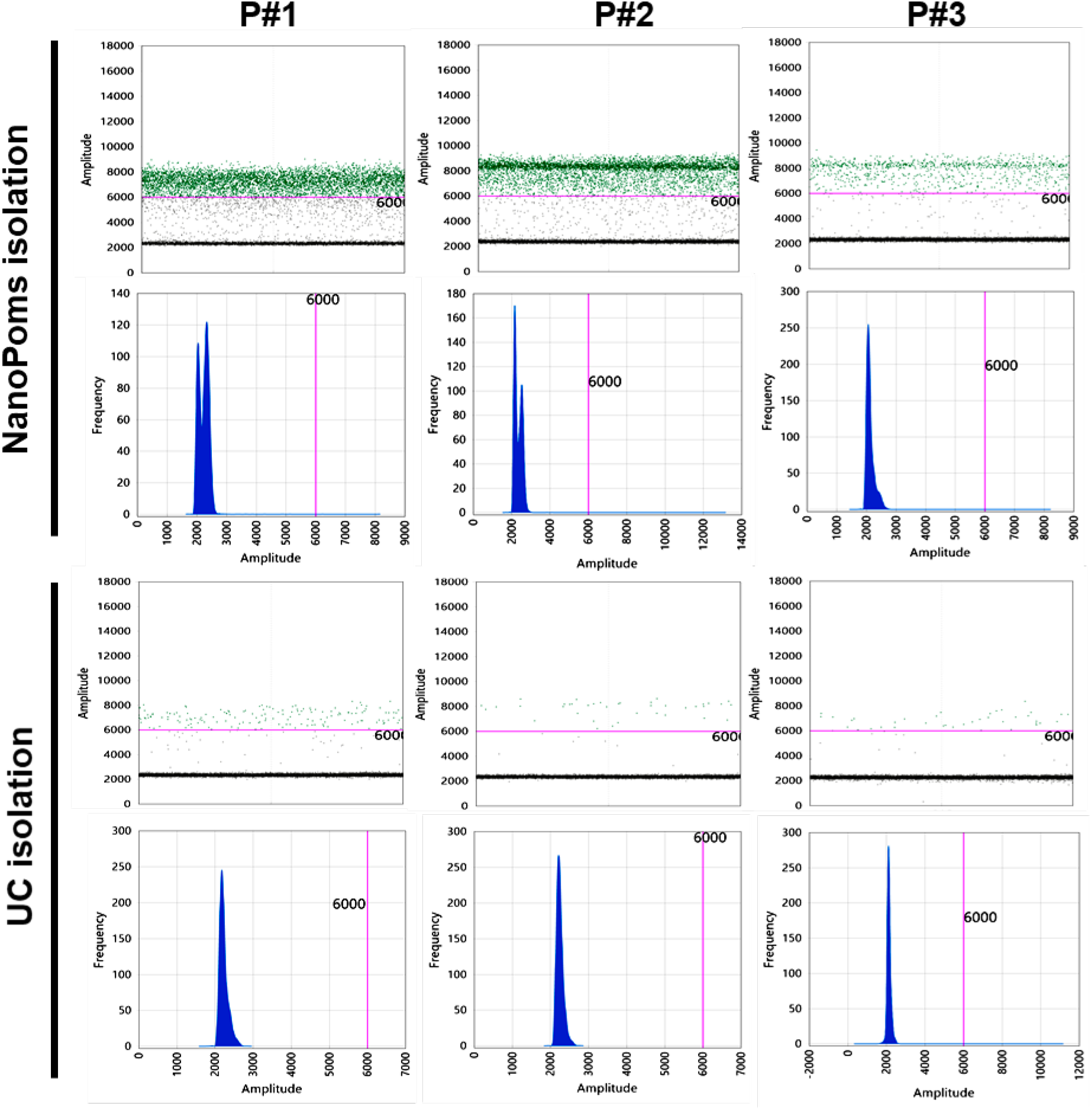
The droplet digital PCR analysis of NanoPoms prepared sEVs detects the EGFR heterozygosity. The EGFR heterozygosity from three bladder cancer patient urine samples was detected using NanoPoms sEVs preparation. In contrast, UC isolated EV DNAs from the same sample input did not lead to the detection of EGFR heterozygosity in patient 2 and patient 3. The DNA copy numbers were substantially lower in UC prepared EV DNAs as compared to NanoPoms.

**Fig. S6.**
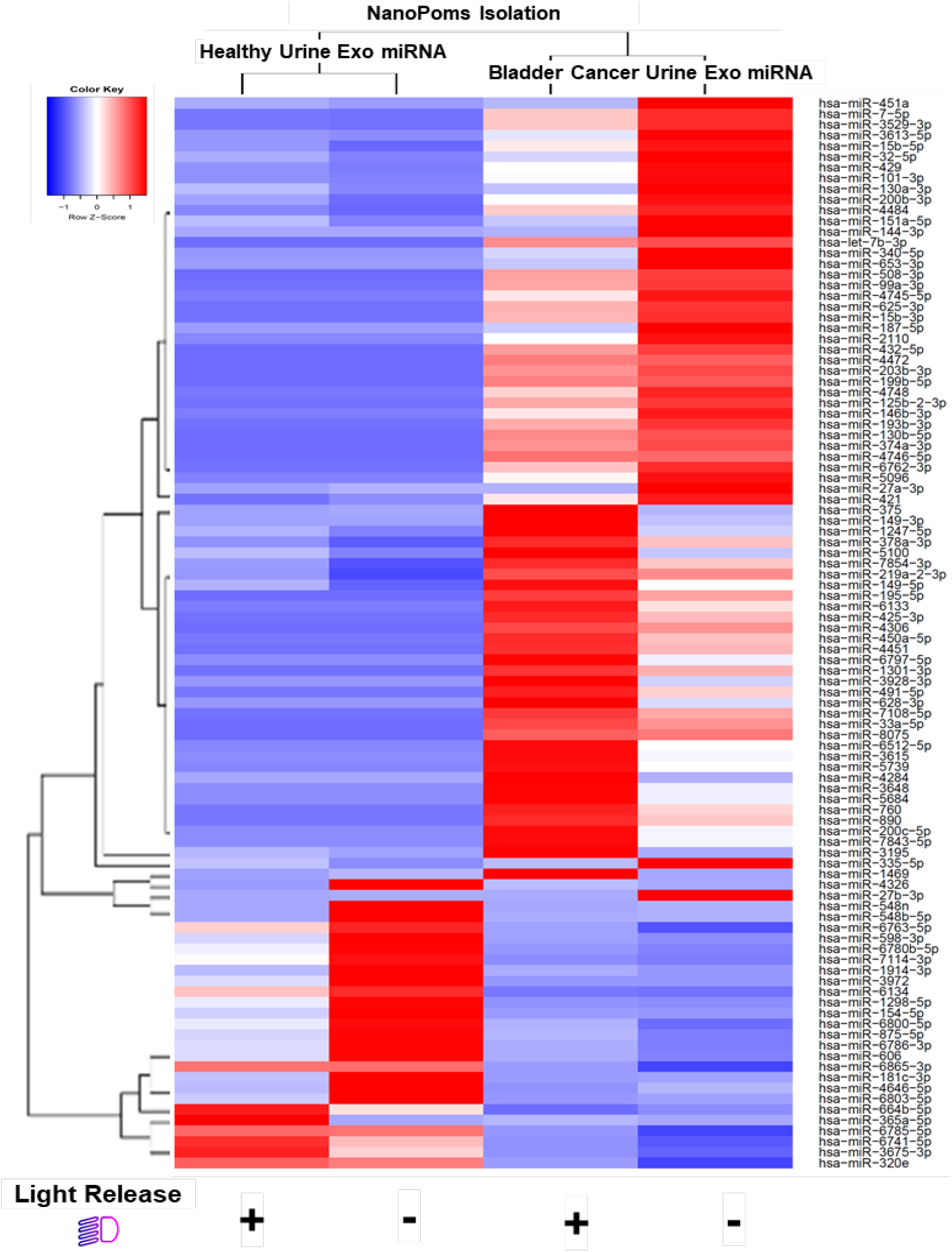
Heatmap dendrogram clustering analysis depicts the top 100 highly expressed miRNAs from urinary sEVs derived both the BC patient and the healthy control. The small RNAs were extracted from NanoPoms sEVs preparation with and without light release process. Red color indicates a higher expression z-score. Hierarchical clustering was performed, using the Spearman correlation method. The disease group can be significantly differentiated from healthy group regardless weather the light release process was implemented.

**Table S1.** Comparative analysis of cost, time, steps, and performance between EV isolation methods.

| **EV Isolation Methods** | **NanoPoms** | **Ultracentrifugation** | **Chromatographic Column** | **Polymer Precipitation** |
| --- | --- | --- | --- | --- |
| **Cost** | Low, no instrument required | High, very expensive equipment cost | Low | Low |
| **Protocol duration** | ~4 hrs | ~12 hrs | ~6 hrs | ~6 hrs |
| **Processing capacity** | From ~ μL to L | ~12- 500 mL | ~ 150 μL-100 mL (IZON) | ~ ≤ 1 mL ExoQuick™ |
| **Subtype specificity** | Surface marker defined specificity to subtypes | No | No | No |
| **Purity** | High to homogeneous subpopulations | Mixture of EV populations | Mixture of EV populations | Mixture of EV with protein aggregates |
| **Reproducibility** | High | Low | - | - |
| **Scalability** | Yes | No | Yes | No |

### NGS analysis of urinary EV small RNAs

#### Bioinformatics Analysis

The sequences of precursor miRNA, tRNA, snRNA, snoRNA, scaRNA, rRNA, scRNA, vaultRNA, lncRNA, miscRNA, pseudogenes, retained introns, ribozymes and transcribed unprocessed pseudogenes were extracted from RefSeq (hg38) and GENCODE (v29) to build a customized database. We refer to this database as the customized ncRNA database. For further quantifying mature miRNA abundance, their sequences were extracted from miRBase1 (ver. hg38) to form another database, which is refer to as the mature miRNA database. Initial quality assessment of the reads was performed using the FASTQC (https://www.bioinformatics.babraham.ac.uk/projects/fastqc/) package. Cutadapt was used to trim the 5’ (GUUCAGAGUUCUACAGUCCGACGAUC) and 3’ (AACTGTAGGCACCATCAAT) adaptor sequences from the sequences. The unique molecular indices (UMI) sequences in the QIAseq libraries were further trimmed using FASTP (--umi_loc=read1 --umi_len=12). The trimmed reads were filtered with a minimum length of 15nt. Mapping to the mature miRNA database and customized non-ncRNA database was performed using BWA (bwa-aln)2. We relaxed the BWA parameter to allow each read to mapped to at most 100 locations, accounting for the numerous multi-mapping cases induced by the short-read length of the sequencing reads. To ensure maximum accuracy, we further restrict perfect sequence in seeding (by setting -k=0). All other parameters were used as default. With an in-house script, we counted the read mapping to quantify the expression level of each ncRNA gene and the abundance of each mature miRNA. For multi-mapping, we evenly distributed its abundances to all mapped locations. The mapping results against the customized ncRNA database were used to quantify different types of ncRNAs as the Pie charts (refer pie-chart figures here). The mapping results against the mature miRNA database were summarized as the volcano plot (refer volcano plot here). The mapping results were further analyzed using the Deseq2 3 to reveal significantly different abundant miRNAs. The most significant 100 miRNAs were further selected to generate the heatmap (shown below).

**Table s2.**
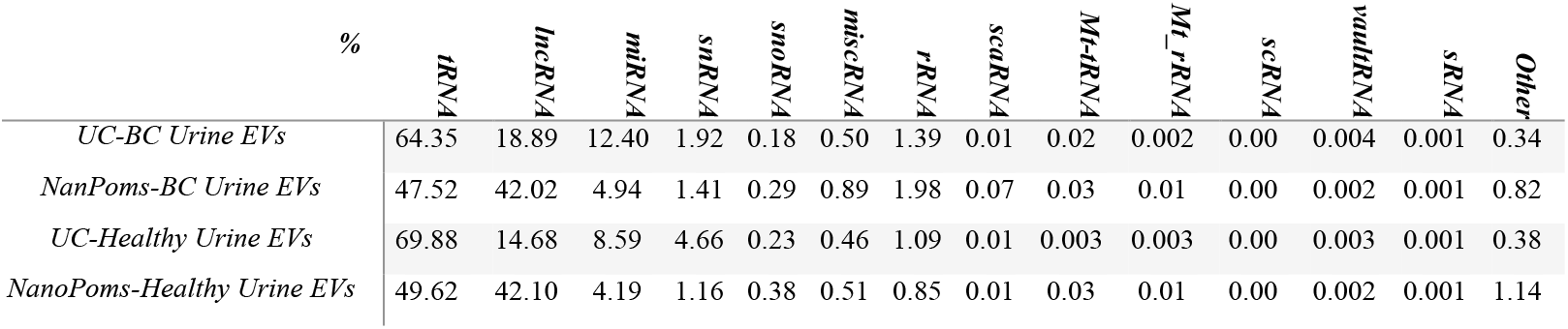
The distribution of small RNAs from urinary EV isolated by UC and NanoPoms.

**Table s3.** The top 10 highly enriched miRNAs identified from NanoPoms isolated urinary sEVs as compared with UC.

| *Genes* | *Log2 Fold Change* | *p value* | *Reported signaling pathway/functions* | *Ref* |
| --- | --- | --- | --- | --- |
| *hsa-miR-3168* | 5.140358266 | 1.38E-07 | KRAS-dependent sorting to exosomes from colorectal cancer cell lines<br>Known Melanoma mature miRNA | 4, 5 |
| *hsa-miR-92b-5p* | 4.358349687 | 1.08E-06 | Cancer metastasis<br>Promote EMT in bladder cancer migration | 6, 7 |
| *hsa-miR-891a-5p* | -5.329381923 | 1.19E-06 | Prognostic marker for HR-positive breast cancer | 8 |
| *hsa-miR-6785-5p* | 5.063745282 | 1.80E-06 | miRNA target genes in the TP53 signaling pathway in tumor | 9 |
| *hsa-miR-934* | -4.078662275 | 3.10E-06 | Cancer metastasis, exosomal oncogene<br>Non-coding RNA with neurogenic function | 10-12 |
| *hsa-miR-6883-3p* | 4.420682437 | 5.64E-06 | Target CDK4/6 in colon cancer cells | 13 |
| *hsa-miR-3202* | 5.039882582 | 8.31E-06 | Regulating TLR signaling pathway<br>Promoted Cell Apoptosis | 14, 15 |
| *hsa-miR-3648* | -4.682609314 | 1.30E-05 | Promote invasion and metastasis of human bladder cancer<br>Regulate cell proliferation | 16, 17 |
| *hsa-miR-6802-5p* | 4.168335059 | 1.84E-05 | Exosomal regulatory miRANs<br>Heart disease | 18, 19 |
| *hsa-miR-6763-5p* | 4.0778099 | 2.21E-05 | Immunity Regulation | 20, 21 |

#### Proteomic analysis of urinary sEV proteins

Urinary EV pellets resultant from ∼2 mL of urine from both bladder cancer patients and healthy individuals were reconstituted in 400 µL of M-PER Mammalian Protein Extraction Buffer (Thermo) supplemented with 1× Halt Protease Inhibitors (Thermo) and sonicated in an ultrasonic water bath for 15 min. Lysates were exchanged into ∼40 µL of 100 mM triethylammonium bicarbonate using Amicon Ultra-0.5, 3 k columns (Millipore). Lysate were digested overnight with Trypsin Gold, Mass Spectrometry Grade (Promega). Peptides were finally reconstituted into 0.1% formic acid to a concentration of 0.1 µg/µL and injected into a 1260 Infinity nHPLC (Agilent) with separation from a Jupiter C-18 column, 300 Å, 5 μm, Phenomenex) in line with a LTQ XL ion trap mass spectrometer equipped with a nano-electrospray source (Thermo). All fragmentation data were collected in CID mode. The nHPLC was configured with binary mobile phases that included 10 min at 5% of 0.1% formic acid and 85% acetonitrile, 180 min (LTQ XL), 5 min wash using 70% of 0.1% formic acid, 85% acetonitrile, and 10 min equilibrate. Samples were performed in duplicate for obtaining the average values utilized for analysis. Searches were performed with UniRef100 database which includes common contaminants from digestion enzymes and human keratins. Peptides were filtered and quantified using ProteoIQ (Premierbiosoft, Palo Alto, CA).

**Table s4.**
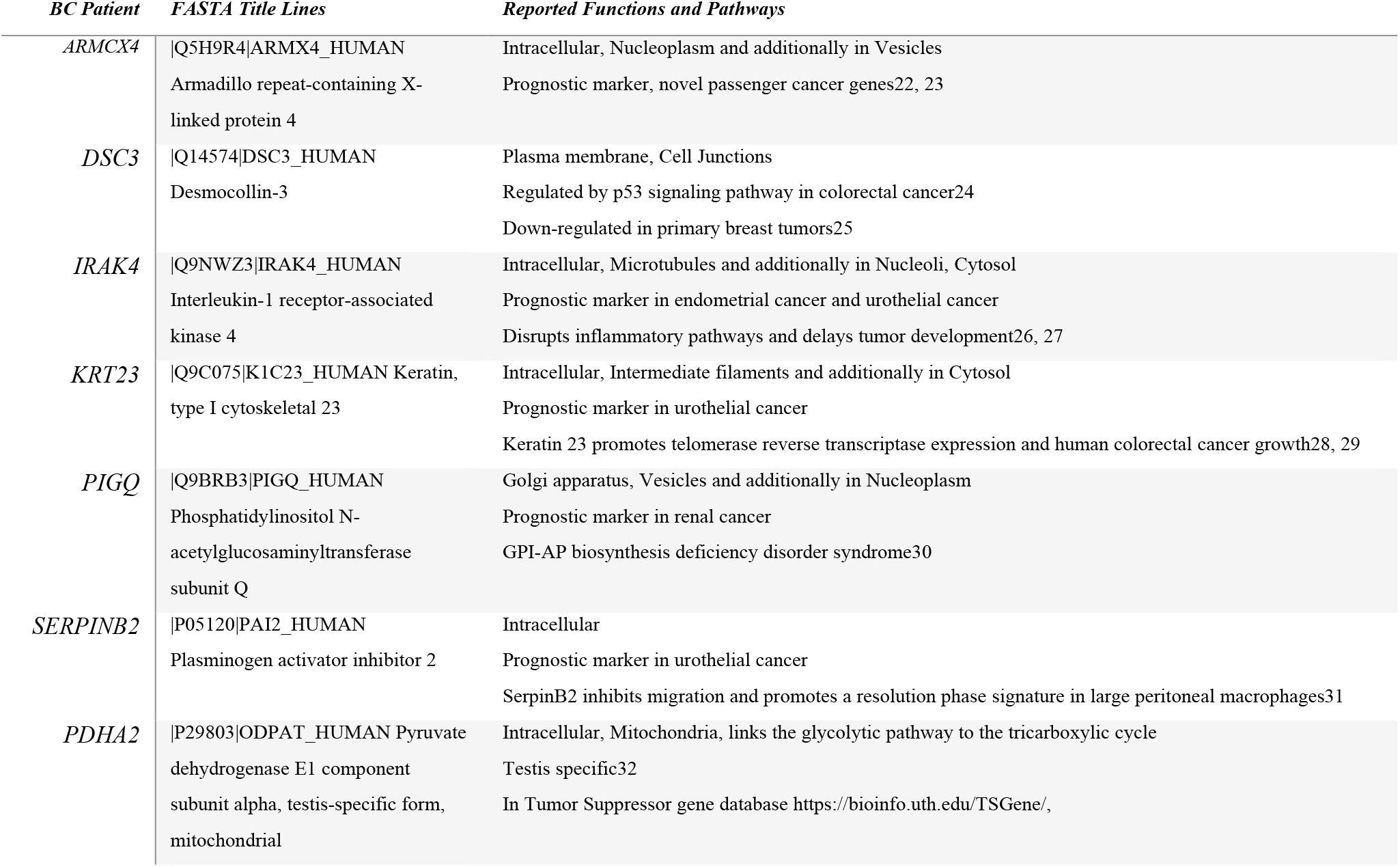
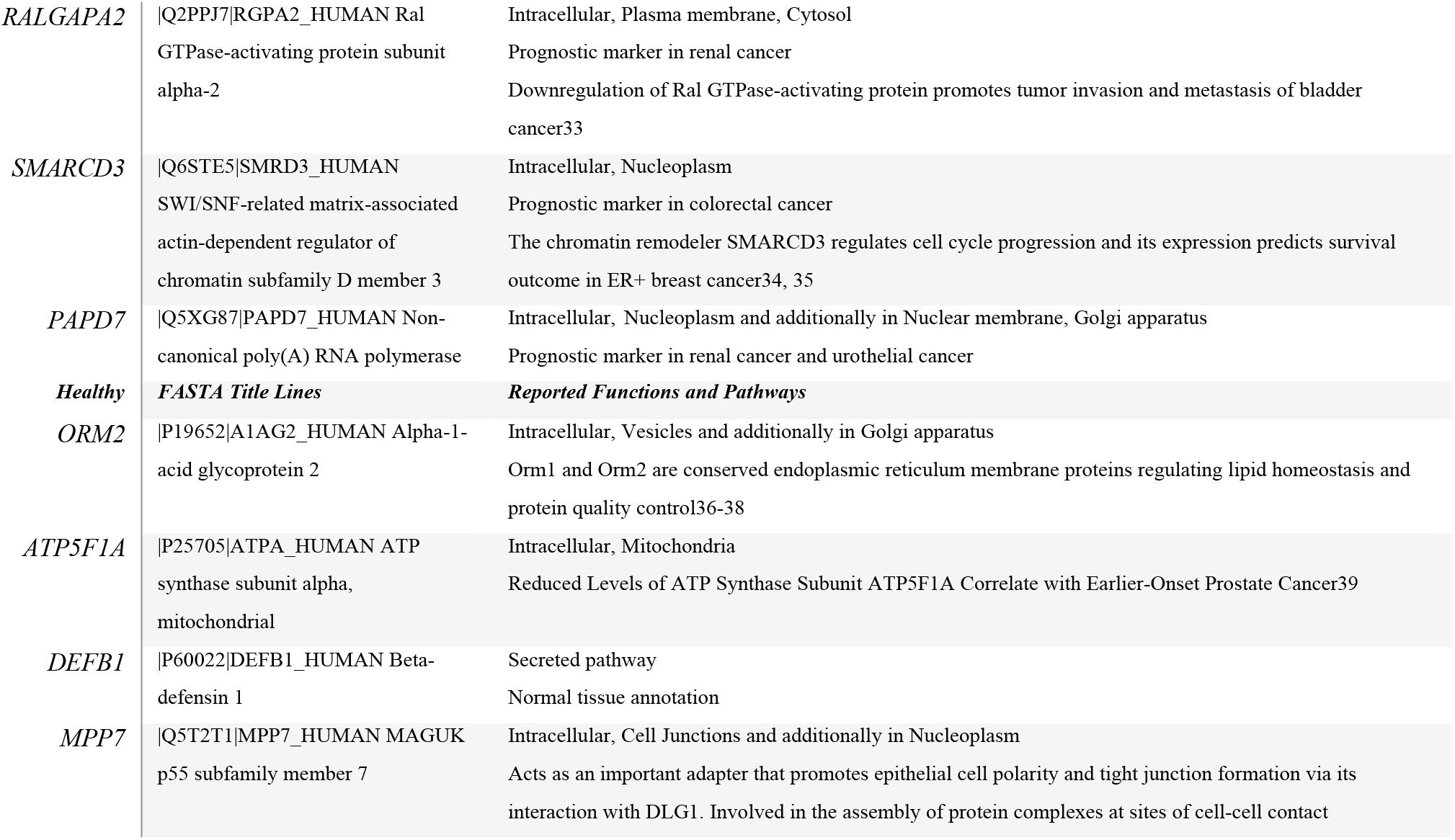
The 10 unique gene products identified from BC patient only and 4 unique genes identified from healthy control group only, by proteomic analysis of NanoPoms isolated urinary sEV proteins. The Human Protein Atlas database was used: https://www.proteinatlas.org/

